# *Dlx5* and *Dlx6* expression in GABAergic neurons controls behavior, metabolism, healthy aging and lifespan

**DOI:** 10.1101/583708

**Authors:** Camille de Lombares, Eglantine Heude, Gladys Alfama, Anastasia Fontaine, Rim Hassouna, Cécile Vernochet, Fabrice de Chaumont, Christophe Olivo-Marin, Elodie Ey, Sébastien Parnaudeau, François Tronche, Thomas Bourgeron, Serge Luquet, Giovanni Levi, Nicolas Narboux-Nême

## Abstract

*Dlx5* and *Dlx6* encode two homeobox transcription factors expressed by developing and mature GABAergic interneurons. During development *Dlx5/6* are important for the differentiation of *Parvalbumin* (*Pvalb*)-expressing neurons. Perinatal lethality of homozygous mice in which *Dlx5/6* have been constitutively deleted has, so far, hindered the study of the function of these genes in adult neurons. We first show that *Dlx5* and *Dlx6* are expressed by all subclasses of adult cortical GABAergic neurons. Then we analyse *Vgat*^*ΔDlx5-6*^ mice in which *Dlx5* and *Dlx6* are simultaneously inactivated in all GABAergic interneurons. *Vgat*^*ΔDlx5-6*^ mice present a behavioral pattern suggesting reduction of anxiety and obsessive-compulsive activities. They rapidly access and spend more time in the central region of an open field, bury few marbles in the marble burying test and show little interest in nest building. Male and female 20-month-old *Vgat*^*ΔDlx5-6*^ animals have the same size as their normal littermates, but present a 25% body weight reduction associated with a marked decline in white and brown adipose tissue. Remarkably, both *Vgat*^*ΔDlx5-6/+*^ and *Vgat*^*ΔDlx5-6*^ mice present a 33% longer median survival than their control littermates. Hallmarks of biological aging such as motility, adipose deposition and coat conditions are improved in mutant animals. Our data imply that GABAergic interneurons can regulate mammalian healthspan and lifespan through *Dlx5/6*-dependent mechanisms. Understanding these regulations can be an entry point to unravel the processes through which the brain affects body homeostasis and, ultimately, longevity and healthy aging.

**SIGNIFICANCE STATEMENT:** Dlx5 and Dlx6 are transcription factors controlling several developmental processes, including GABAergic neuronal migration and differentiation. To study their function in adult brain, we selectively inactivated both genes in GABAergic interneurons (*Vgat*^*ΔDlx5-6*^ mice). Mutant mice have reduced anxiety-like and obsessive-compulsive behaviors. Interestingly, *Vgat*^*ΔDlx5-6*^ mice have a 25% body weight reduction and about 70% less white and brown adipose tissue; their general health status is excellent. *Vgat*^*ΔDlx5-6*^ mice have a median survival about 33% longer than their control littermates and hallmarks of biological aging are improved. Dlx5/6-dependent regulations in GABAergic neurons could be an entry point to understand how the brain determines the psychophysiological status of the body and, ultimately, longevity and healthy aging.

## INTRODUCTION

Brain activity depends on GABAergic inhibitory interneurons, an heterogeneous class of neurons distinguished by diverse anatomical, biochemical and physiological characteristics (1). GABAergic interneurons are found in the cerebral cortex (2) and in most other brain regions, including the hypothalamus (3); their activity affects functions as diverse as cognition, pain transmission (4) and feeding behavior (5). Classically, more than 20 categories of inhibitory GABAergic interneurons have been described in the cortex and the hippocampus (2, 6), however, the extent of GABAergic cellular diversity begins only recently to be appreciated thanks to single cell transcriptomic analysis (7-10). To generate these diverse morphotypes, neuronal progenitors engage in stereotyped transcriptional trajectories in which combinatorial sequences of transcription factors (TFs) progressively unfold specific differentiation programs (11-13). During mouse development, cortical interneurons derive from progenitors located in the embryonic subcortical telencephalon (subpallium) (13-15). In particular, the majority of Parvalbumin (Pvalb) and Somatostatin (Sst) positive GABAergic interneurons originate in the medial ganglionic eminence (MGE), one of the morphological subdivisions of the subpallium (16, 17), and reach their final destination in the mature brain via tangential migration (18).

*Dlx* genes encode a family of homeodomain transcription factors that control multiple aspects of embryonic development (19) including neurogenesis (20). In mammals, six *Dlx* genes are arranged in three pairs of closely linked transcription units (21, 22). *Dlx1, Dlx2, Dlx5*, and *Dlx6* are expressed in precursors of the GABAergic lineage (23). During early development, *Dlx5* is initially expressed in the anterior neural ridge and its derivatives (24). At later stages, *Dlx* expression follows a temporal, positional, and functional sequence in the ventricular/subventricular (VZ/SVZ) zone of the embryonic ganglionic eminence (GE) (25, 26): *Dlx2* and *Dlx1* are mainly found in neuroepithelial cells of the VZ, while *Dlx5* is mostly expressed in cells of the SVZ. Later in embryogenesis, *Dlx5* is expressed by cells of the rostral migratory stream (RMS), and of the olfactory bulb (OB) (20, 25, 27, 28).

In the adult brain, the levels of *Dlx5* expression diminish considerably, however, analysis of the expression of a *Dlx5/6* enhancer/reporter construct (29) and of a *Dlx5* BAC (14) in transgenic mice suggests that a low expression level is maintained in mature GABAergic interneurons.

The function of Dlx5/6 in adult GABAergic neurons has been, so far, difficult to analyze due to perinatal lethality of mutant mice (30-33). Nonetheless, grafting of immature mutant interneurons into wild type newborn brains results in the reduction of Pvalb-positive GABAergic neurons in the adult (14). The importance of *Dlx* genes for GABAergic differentiation has been further demonstrated by forcing or inhibiting expression of *Dlx2* or *Dlx5* in cortical neurons (20, 34, 35). Dlx5 promotes GABAergic differentiation by participating to complexes containing MAGE-D1 and Necdin (36, 37). Interestingly, loss of *Necdin* gene expression is associated with Prader-Willi Syndrome (PWS) (38), a neurobehavioral disorder characterized by hyperphagia and mental health disorders with accelerated aging (39).

In humans, *DLX5* is located on chromosome 7q21.3 and is part of a gene cluster imprinted in lymphoblasts and brain tissues (40). In the mouse brain, however, *Dlx5* is biallelically expressed with preferential transcription of the maternal allele (41, 42). An interesting association between *DLX5* and the aging process comes from the linear correlation observed between aging and hypermethylation of *DLX5* (43, 44) or during senescence of human mesenchymal stem cells (45, 46). It has been shown that MECP2 binds to the *Dlx5/6* locus suggesting its possible implication in Rett syndrome (42, 47).

Although *DLX5* and *DLX6* are important for the development of cortical GABAergic interneurons (48) their distribution and function in the adult brain (29) and their implication in neuropsychiatric conditions remain elusive. Preadolescent mice, heterozygous for a generalized deletion of *Dlx5* and *Dlx6* (*Dlx5/6*^+/-^) (49), present traits reminiscent of human schizophrenia, but also gonadal (50), bone (51) and craniofacial anomalies (52) not associated to GABAergic interneurons.

Here we analyse the phenotype of *Vgat*^*ΔDlx5-6*^ mice in which *Dlx5* and *Dlx6* are both inactivated only in GABAergic interneurons. Heterozygous and homozygous mutants (*Vgat*^*ΔDlx5-6/+*^ and *Vgat*^*ΔDlx5-6*^) present a reduction in anxiety-like and obsessive-compulsive-like behaviors, have less adipose tissue and live 33% longer and in better health than their littermates. We conclude that Dlx5/6-dependent regulations in GABAergic interneurons affect behavior as well as metabolism, and contribute to determine heathspan and lifespan.

## RESULTS

### Inactivation of *Dlx5/6* in adult GABAergic neurons

We first analysed *Dlx5*^*lacZ/+*^ mice in which *Dlx5* exons I and II are replaced by the *E. coli lacZ* gene and β-galactosidase activity reproduces the known pattern of expression of the gene in embryonic (30) and adult tissues (53). In the central nervous system (CNS) of adult *Dlx5*^*lacZ/+*^ mice, β-galactosidase activity is widely expressed in forebrain regions including the cerebral cortex, the striatum and the hypothalamus (Fig. 1A-C). Double immunofluorescence labelling showed that most cortical Parvalbumin (84%), Calretinin (100%) and Somatostatin (89%) interneurons are positive for *Dlx5* (Fig. 1D-D”). Single cell RNA sequencing analysis (scRNA-seq) of publicly available data sets (9, 54) showed that *Dlx5* and *Dlx6* expression is restricted to all subtypes of GABAergic interneurons characterized by expression of *Gad1/2* and *Vgat* including *Sst, Pvalb, HTr3a, Npy* and *CR* clusters (Figs. 1E, S1). In contrast, *Dlx5/6* expression was not detected in glutamatergic neurons (Fig. S1, *Vglut2*) nor in mature astrocytes (*Aldh1l1*) and oligodendrocytes (*Olig2*) (Fig. S1).

**Figure 1).**
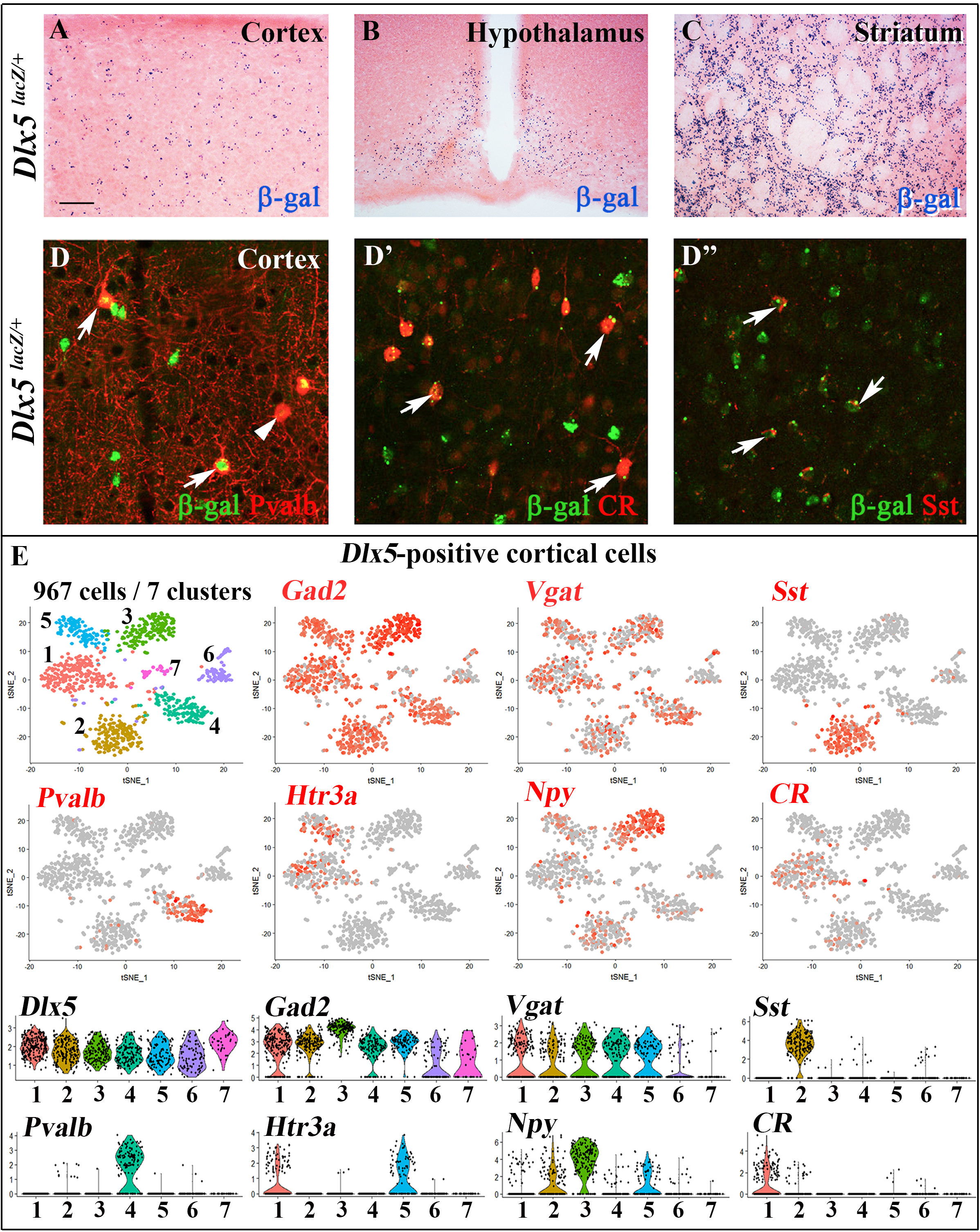
Expression of *Dlx5* in adult brain. A-C) Sections from adult brain of *Dlx5*^*lacZ/+*^ mice. β-D-galactosidase activity, visualized as dark blue dots, is evident in the cortex (A), hypothalamus (B) and striatum (C). D-D’’) Adult brain somatosensory cortex from *Dlx5*^*lacZ/+*^ mice was double stained with anti-β-D-galactosidase antibodies (green) and antibodies against major GABAergic neuronal subclasses (Parvalbumin (Pvalb), Calretinin (CR) and Somatostatin (Sst)) (red). Arrows point to examples of double-stained neurons; arrowhead indicates a β-D-galactosidase-negative/Pvalb-positive neuron. Bar: 250 μm A-C; 25 μm D, D’’. E) (Upper panels) t-distributed stochastic neighbor embedding (t-SNE) plots showing the relationship among 967 *Dlx5*-positive single cells isolated from the frontal cortex using principal component analysis (PCA). The seven resulting cell clusters are color-coded and distribution of selected markers for different classes of cortical GABAergic neurons are presented (*Gad2, Vgat, Sst, Pvalb, Htr3a, Npy, CR*). All *Dlx5*-positive clusters are *Gad2* and *Vgat*-positive, all major GABAergic subtypes include *Dlx5*-positive neurons. (Lower panels) Violin plots showing the expression of the selected marker among the *Dlx5*-positive subclusters. Gene expression level is presented on linear scale and adjusted for different genes.

To inactivate *Dlx5*/*6* genes in GABAergic interneurons we crossed *Dlx5/6*^*flox/flox*^ mice, in which the homeodomain-encoding regions of both *Dlx5* and *Dlx6* are flanked by *lox* sequences (53) with *Vgat*^*cre/+*^ mice in which an IRES-Cre recombinase cassette is inserted downstream of the stop codon of the endogenous *Vgat* (vesicular GABA transporter) gene. In *Vgat-cre* mice *Cre-recombinase* expression is observed in all GABAergic neurons, but not in other cell types (55). We thus generated *Dlx5/6*^*flox/+*^;*Vgat*^*Cre/+*^ (thereafter designated *Vgat*^*ΔDlx5-6/+*^) mice which, when crossed with *Dlx5/6*^*flox/flox*^ mice, generated *Vgat*^*ΔDlx5-6*^ individuals (*Dlx5/6*^*flox/flox*^;*Vgat*^*Cre/+*^) (Fig. S2A). *Vgat*^*ΔDlx5-6/+*^ and *Vgat*^*ΔDlx5-6*^ mice were viable and fertile. We choose to inactivate both *Dlx5* and *Dlx6* in GABAergic interneurons since these two closely related genes have often redundant functions (31, 32). PCR analyses of cortical DNA confirmed that recombination had occurred in *Vgat*^*ΔDlx5-6/+*^ and *Vgat*^*ΔDlx5-6*^ mice (Fig. S3B). RT-PCR confirmed the absence of exon II in the vast majority of *Dlx5* cortical transcripts (compare second and first lane of Fig. S3B), and in all *Dlx6* cortical transcripts of *Vgat*^*ΔDlx5-6*^ mice (Fig. S3A-B).

### Behavioral defects associated with *Dlx5/6* inactivation in GABAergic neurons

To examine how *Dlx5/6* inactivation in GABAergic neurons affects adult mouse behavior we scored the performance of heterozygous and homozygous mutant mice in five different tests: locomotor activity, open field, marble burying, nest building and socialization.

#### Open Field Test (OFT)

Mice were filmed while freely moving in a 72×72 cm square flat arena for 10 minutes; the path followed by the animals was tracked and analysed (Fig. 2A). Vgat^*ΔDlx5-6/+*^ and *Vgat*^*ΔDlx5-6*^ mice travelled respectively two or four times more the distance than their control littermates (Fig. 2A, B). The peak and mean velocity reached by *Vgat*^*ΔDlx5-6/+*^ and *Vgat*^*ΔDlx5-6*^ mice was 2 to 4 times higher than that of their control littermates (Fig. S4A, B) moreover, the acceleration of *Vgat*^*ΔDlx5-6*^ mice was up to six times higher than control and three times higher than heterozygous mutants (Fig. S4C).

**Figure 2).**
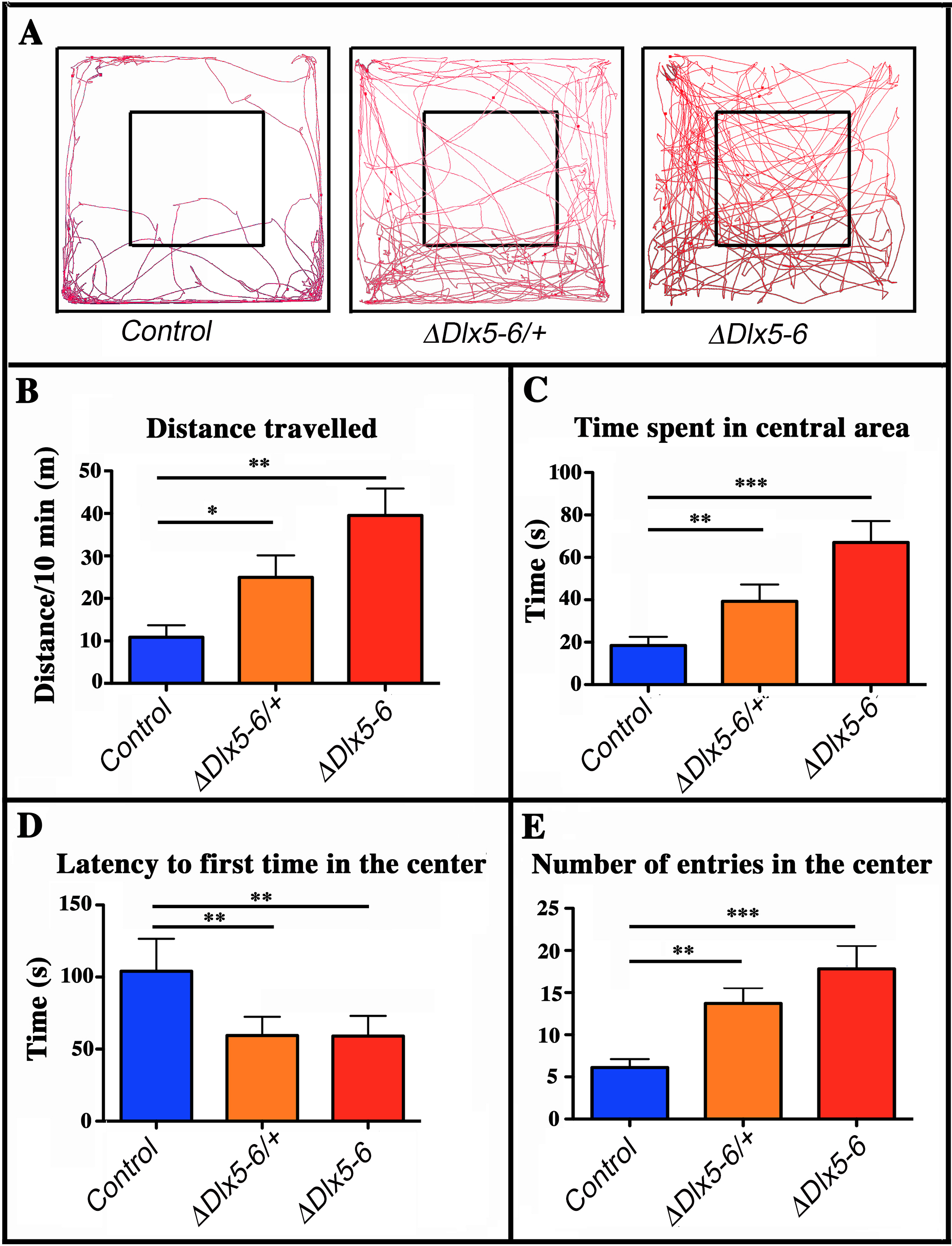
Behavioral response in an open field test. A) Travel pathway (red) of illustrative examples of control, *Vgat*^*ΔDlx5-6/+*^ and *Vgat*^*ΔDlx5-6*^ mice *(ΔDlx5-6/+* and *ΔDlx5-6* respectively) filmed for 10 min in the open field test arena (72×72 cm larger square perimeter). The central region (36×36 cm) is indicated by the smaller square. *Vgat*^*ΔDlx5-6/+*^ and *Vgat*^*ΔDlx5-6*^ animals travelled a significantly longer distance than their control littermates (B) and spent more time in the center of the open field (C), where they entered more rapidly (D) and more frequently (E). Histograms bars indicate the mean±SEM; One way ANOVA, post hoc analysis by Tukey’s HSD (Controls: n = 45 and *Vgat*^*ΔDlx5-6/+*^: n = 38, *Vgat*^*ΔDlx5-6*^ : n=34): ***: p<0.001; **: p<0.01; *: p<0.05.

The time spent in the center of the open field was significantly increased in both *Vgat*^*ΔDlx5-6/+*^ and *Vgat*^*ΔDlx5-6*^ animals (Fig. 2C). Both heterozygous and homozygous mice entered in the central area of the open field faster (50% reduction in latency) (Fig. 2D) and more frequently (Fig. 2E) than control littermates. Control mice spent more time in the corners of the open field than mutants (Fig. 2A). In the OFT no difference was observed between males and females. Globally, the findings of the OFT suggest that adult mutants present a reduction in anxiety-like behavior.

#### Locomotor activity

Locomotor activity during a 90 min period was measured for control and *Vgat*^*ΔDlx5-6*^ mice for 3 consecutive days. Control mice displayed normal spontaneous locomotor response and habituation to a novel environment with a high locomotor activity on day 1 that then decreased and stabilized on days 2 and 3. In contrast, *Vgat*^*ΔDlx5-6*^ mice showed a significantly higher locomotor response on day 1 compared to controls; however on days 2 and 3 no significant difference was detected (Fig. S4D). Altogether these results suggest that *Vgat*^*ΔDlx5-6*^ mice do not show a generalized hyperactivity, but are more reactive to novelty.

#### Marble burying test (MBT)

The consequences of *Dlx5/6* inactivation in GABAergic neurons on stereotyped repetitive behavior were assessed through the Marble Burying Test (MBT) (Fig. 3A).

**Figure 3).**
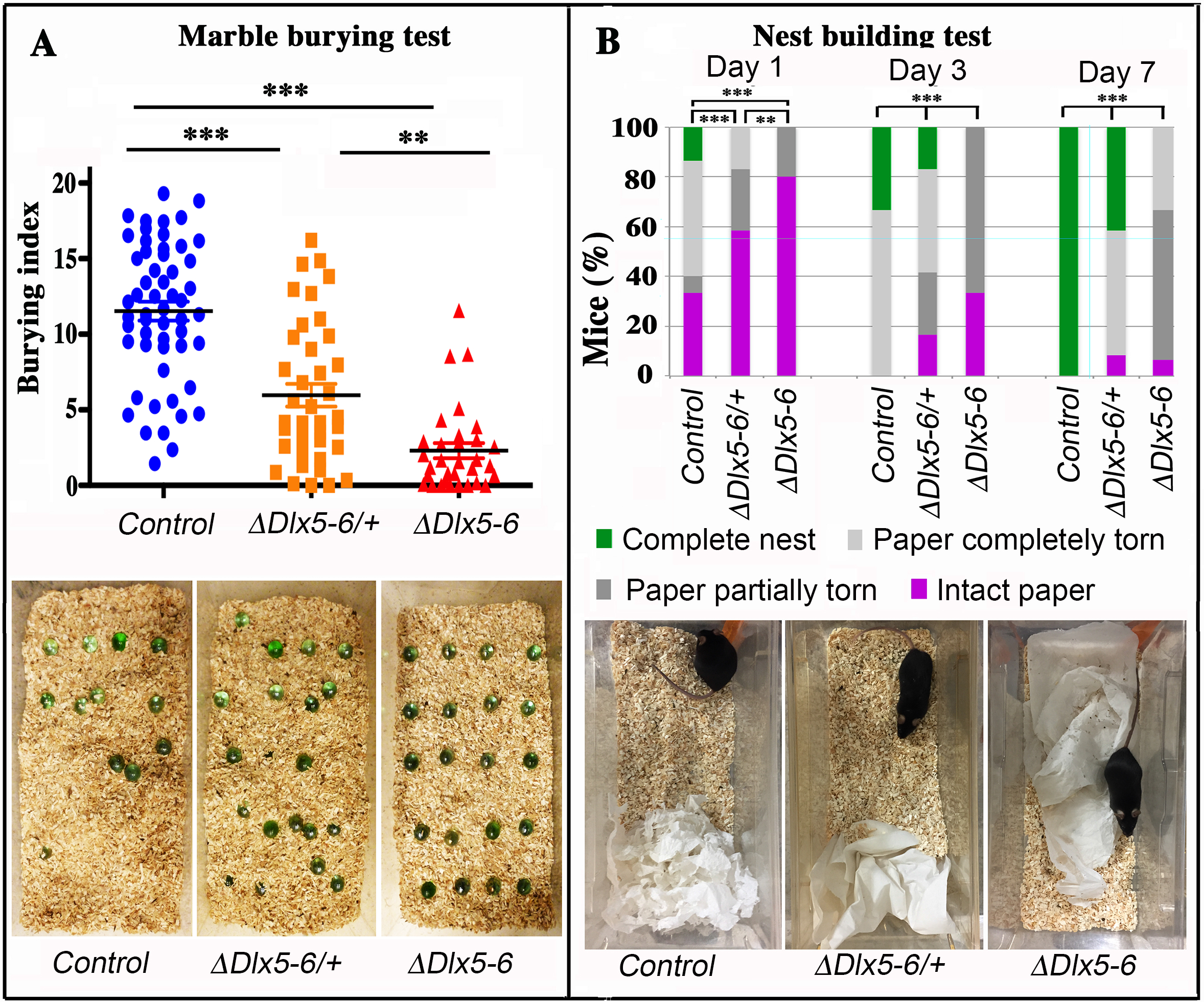
Behavioral response to marble burying and nest building tests. A) The number of marbles buried by each mouse over a 10 min period (Burying index) is plotted to show the dispersion of the results and to highlight the proportion of mutant mice that did not bury even one marble during the test. Horizontal lines indicate median values. (Kruskal-Wallis test; Control: n = 55; *Vgat*^*ΔDlx5-6/+*^: n = 39; *Vgat*^*ΔDlx5-6*^: n=31). The lower panel presents representative examples of the MBT arena at the end of the test for the three genotypes. B) The quality of nests built by control, *Vgat*^*ΔDlx5-6/+*^ and *Vgat*^*ΔDlx5-6*^ mice was evaluated each day over a seven days period. The quality of nest building was scored as: completed nest (green), completely torn paper (light grey), partially torn paper (dark grey) or intact paper (purple). The percentage of mice in each category is indicated at 1, 3 and 7 days. The lower panel presents representative examples of the nest-building arena at the end of the first 24 h of test for the three genotypes. At the end of the scoring period all control mice had completed nest building whereas none of the *Vgat*^*ΔDlx5-6*^ mice had done so (Pearson’s chi-squared; Control: n = 15 and *Vgat*^*ΔDlx5-6/+*^ : n = 12, *Vgat*^*ΔDlx5-6*^ : n=15; ***: p<0.001; **: p<0.01).

During the 10 min test, both *Vgat*^*ΔDlx5-6/+*^ and *Vgat*^*ΔDlx5-6*^ animals buried a significantly lower number of marbles than control littermates (Fig. 3A). Remarkably, 42% (13/31) of the *Vgat*^*ΔDlx5-6*^ animals did not bury or displace even one marble while only 7,7% (3/39) of the *Vgat*^*ΔDlx5-6/+*^ animals presented this extreme phenotype which was never observed in the control group. Mutant animals and control littermates moved similarly throughout the cage, however, mutant mice passed over the marbles without stopping to bury them with rapid movements of their hind limbs as did their control littermates.

#### Nest building test

Nest building is an important natural behavior occurring without intervention or presence of the experimenter. We observed a statistically significant difference in the quality of constructed nest among control, *Vgat*^*ΔDlx5-6/+*^ and *Vgat*^*ΔDlx5-6*^ animals. The quality of the nests built by heterozygous and homozygous mutants was significantly lower than that of control mice (Fig 3B). The difference was already evident after one day (Fig. 3B lower panels) and persisted for at least seven days when the observation was terminated. At the end of the test, none of the *Vgat*^*ΔDlx5-6*^ mice had built a high quality nest (Fig. 3B), whereas all control mice had completed nest construction and only 40% *Vgat*^*ΔDlx5-6/+*^ mice had generated high quality nests (Fig. 3B).

#### Sociability tests

The social behavior of *Vgat*^*ΔDlx5-6/+*^ and *Vgat*^*ΔDlx5-6*^ animals was evaluated in two different tests in open field. In the first test, sociability was measured by comparing the time spent in proximity of a prison occupied by an unfamiliar animal to the time spent near a similar empty prison. In this test all animals spent more time in proximity of the occupied prison, however no significant difference could be seen between control, *Vgat*^*ΔDlx5-6/+*^ and *Vgat*^*ΔDlx5-6*^ animals (Fig. S5). In the second test one control mouse was confronted in an open field to a control, a *Vgat*^*ΔDlx5-6/+*^ or a *Vgat*^*ΔDlx5-6*^ second individual. Their interactions were filmed and analysed. No difference was found between genotypes (Fig. S6).

### Metabolic consequences of *Dlx5/6* inactivation in GABAergic neurons

Throughout their life both male and female *Vgat*^*ΔDlx5-6*^ mice had a similar length, but presented a significantly lower body weight compared to control littermates. In most age groups, heterozygous mutants presented a similar, but less pronounced weight reduction (Fig. 4A, Fig. S7). After 5 months of age both female (Fig. 4A) and male (Fig. S7C) *Vgat*^*ΔDlx5-6/+*^ and *Vgat*^*ΔDlx5-6*^ animals presented up to 25% body weight reduction compared to their normal littermates. Body mass reduction was already evident during growth (Fig. S7A,B) and persisted in adult and aging animals (Figs. 4A, S7C). At any age analysed the nose-to-anus length of normal and mutant animals was not significantly different suggesting that the loss in body weight depended on reduce adiposity.

**Figure 4).**
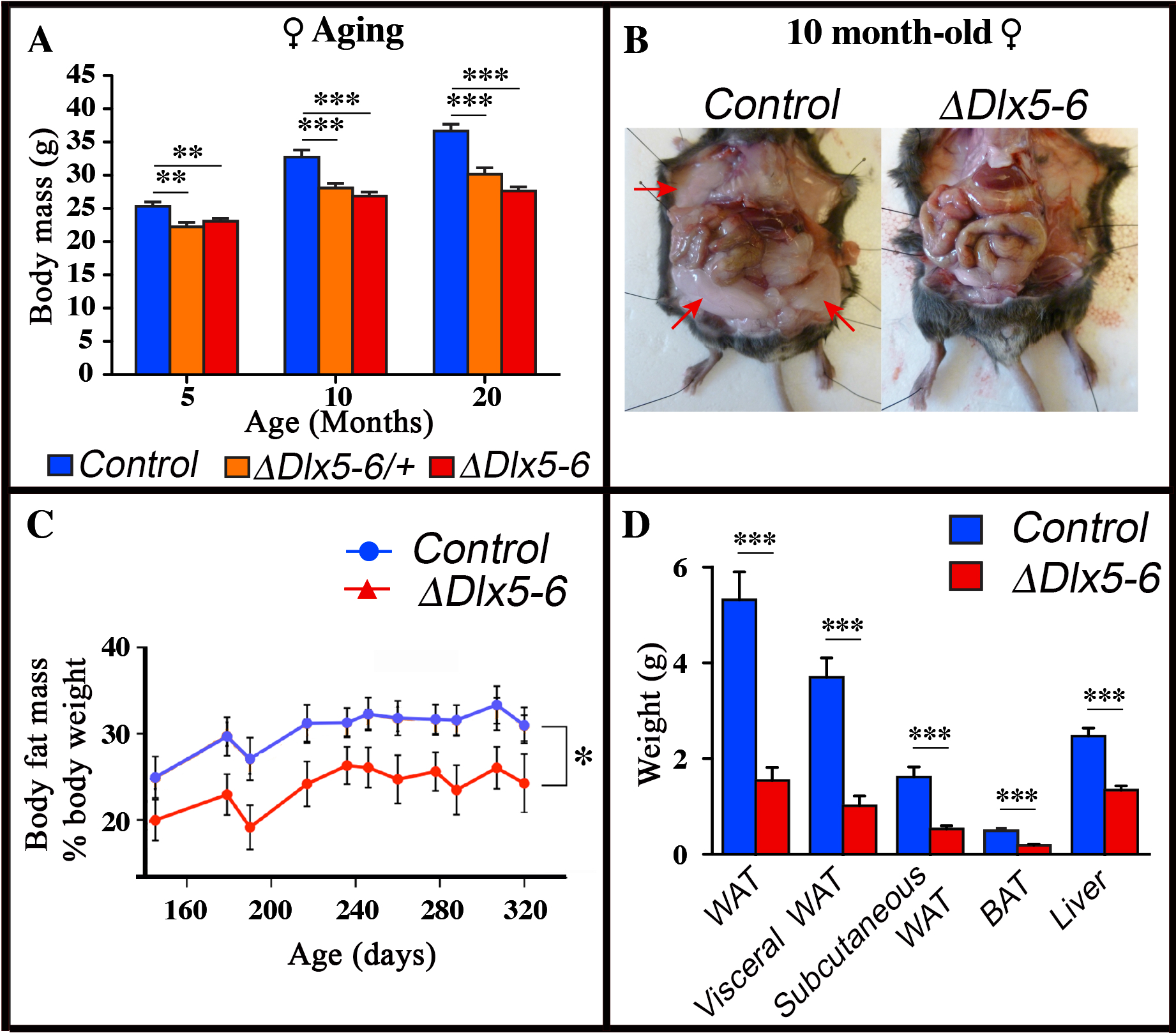
Reduced weight and adipose tissue in *Vgat*^*ΔDlx5-6/+*^ and *Vgat*^*ΔDlx5-6*^ mice. A) The body weight of a cohorts of female control, *Vgat*^*ΔDlx5-6/+*^ and *Vgat*^*ΔDlx5-6*^ mice (n ≥ 8 per group) was measured during the first 20 months of aging. At all time points analyzed *Vgat*^*ΔDlx5-6/+*^ and *Vgat*^*ΔDlx5-6*^ mice (male (Fig. S7) and female) presented a highly significant weight reduction. B-D) Gross anatomical inspection (B), MRI analysis (C) and dissected organ weight (D) confirmed a dramatic reduction of visceral and subcutaneous WAT and of BAT in *Vgat*^*ΔDlx5-6*^ mice (n=11 controls, n=7 *Vgat*^*ΔDlx5-6*^). Mann-Witney test; ***: p<0.001; **: p<0.01; *: p<0.05.

Dissection of 10 months old *Vgat*^*ΔDlx5-6*^ mice (n=7) showed a dramatic reduction (Fig. 4B-D) of visceral (−73% w/w) and subcutaneous White Adipose Tissue (−67% w/w) (vWAT and scWAT) and of Brown Adipose Tissue (BAT) (−62% w/w). Body composition analysis using nuclear magnetic resonance imaging confirmed the reduction in the percent of adipose tissue present in mutant animals (Fig.4C).

### Extended lifespan and healthspan of *Dlx5/6*^*VgatCre*^ mice

Both *Vgat*^*ΔDlx5-6/+*^ and *Vgat*^*ΔDlx5-6*^ mice lived considerably longer than their control littermates. Heterozygous or homozygous inactivation of *Dlx5/6* in GABAergic neurons resulted in prolonging by 33% the median survival of the animals (Fig. 5A) (n=21 controls, 16 *Vgat*^*ΔDlx5-6/+*^ and 17 *Vgat*^*ΔDlx5-6*^). At 18 months, the aging mutant mice appeared in better health than their control littermates. Whereas control mice gained excessive weight at old age, *Vgat*^*ΔDlx5-6/+*^ and *Vgat*^*ΔDlx5-6*^ mice maintained a stable body mass. The external appearance (alopecia; coat conditions; loss of fur color; loss of whiskers) was quantified on two groups (n=12 each) of control and mutant animals at 2, 9 and 18 months of age in order to follow changes in this indicator of aging (56). As shown in Fig. 5B the decline in coat condition of mutant mice was much slower than that observed in their control littermates.

**Figure 5).**
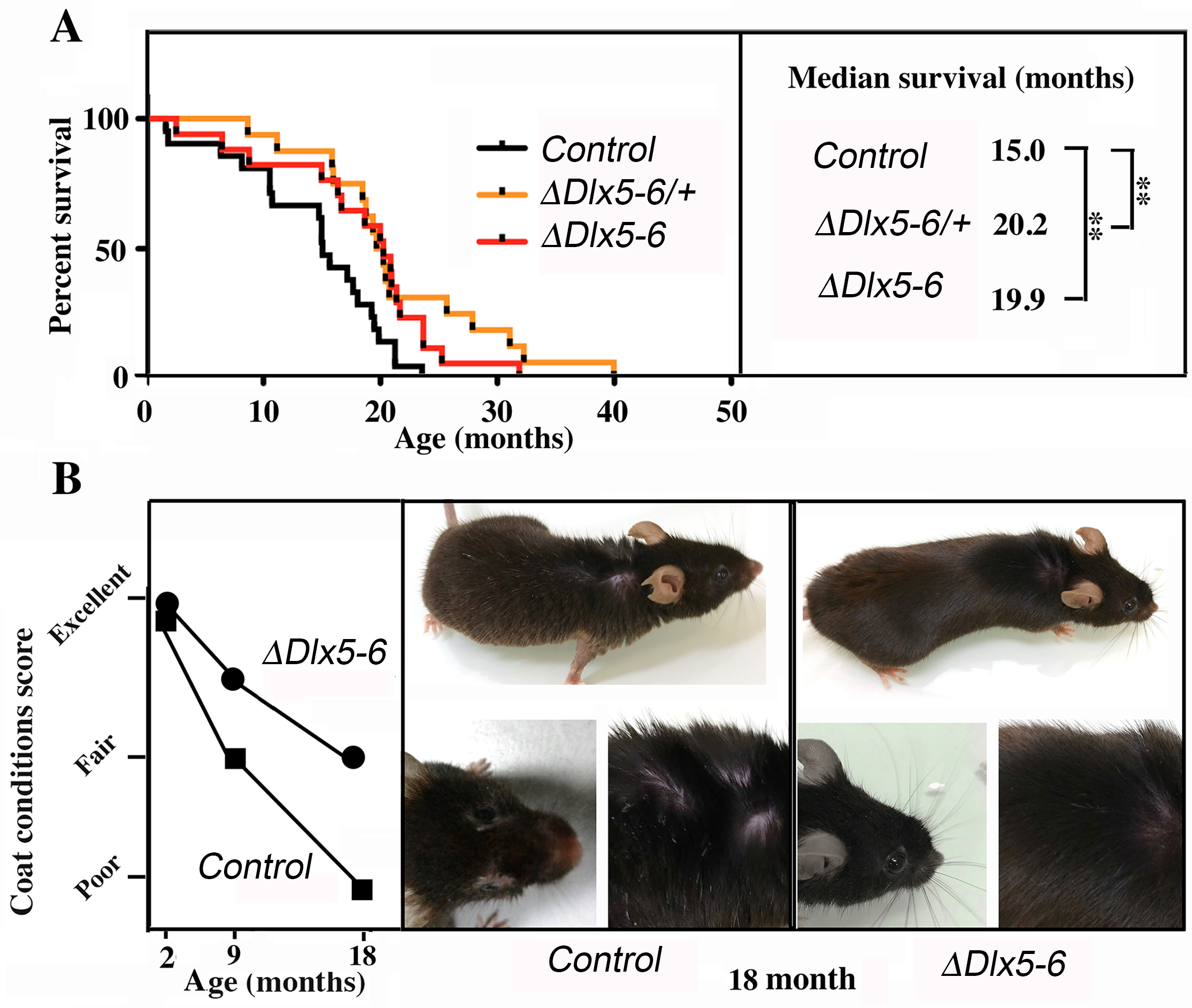
Extended lifespan and healthspan of *Vgat*^*ΔDlx5-6/+*^ and *Vgat*^*ΔDlx5-6*^ mice. A) Kaplan-Meier survival plots. The median survival of *Vgat*^*ΔDlx5-6/+*^ and *Vgat*^*ΔDlx5-6*^ mice is 33% longer than that of their control littermates. B) Scoring of coat conditions of aging control and *Vgat*^*ΔDlx5-6*^ mice. At 2, 9 and 18mo the external appearance (alopecia; coat conditions; loss of fur colour; loss of whiskers) was quantified on two groups (n=12 each) of mutant animals and normal controls (56). Right panels show the coat conditions of two representative control and *Vgat*^*ΔDlx5-6*^ 18mo old animals. Log rank test was performed, **: p<0.01.

These observations, together with the high level of alertness, motility and reduced adiposity of *Vgat*^*ΔDlx5-6/+*^ and *Vgat*^*ΔDlx5-6*^ mice suggest that inactivation of *Dlx5/6* in GABAergic neurons results in a prolonged and healthier life span.

## DISCUSSION

The main finding of this study is that transcriptional modifications limited to GABAergic neurons are sufficient to prolong healthspan and lifespan. Indeed, we have shown that inactivation of the two transcription factors *Dlx5* and *Dlx6* in mouse GABAergic interneurons produces behavioral and metabolic changes accompanied by a prolonged median survival in good health. *Vgat*^*ΔDlx5-6*^ mice are characterized by a reduction in anxiety and compulsive repetitive-like activities, by a remarkable decrease in white and brown adipose tissues and by a 33% median lifespan increase. By single-cell RNAseq analysis and histological techniques, we show that all adult cortical GABAergic neuronal subtypes express *Dlx5* and *Dlx6*. Similar results were obtained analyzing striatum and hypothalamus published datasets (9, 54). All these regions are involved directly or indirectly in the central control of feeding behavior.

The molecular and cellular origin of the phenotypes displayed by *Vgat*^*ΔDlx5-6*^ animals is yet to be deciphered. Interestingly, during development, *Dlx5* expression is sufficient to induce both GABA synthetizing enzymes, *Gad65* and *Gad67* (28, 57). Furthermore, ectopic expression of *Dlx5* in the cerebral cortex can induce *Gad65* expression (29). In humans, alterations of *GAD65/67* have been consistently implicated in cognitive deficits including bipolar disorders and schizophrenia (58, 59). Linkage analysis has identified GAD65 as one of the few genes associated to obesity (60).

Metabolic status is one of the major determinants of healthy aging and life expectancy (61). In turn, metabolism is controlled by the activity of neuronal networks capable to integrate hormonal signals from peripheral organs and cognitive inputs from the central nervous system (62, 63). Therefore, specific brain circuits can integrate sensory, cognitive and physiological inputs and affect the psychophysiological status of the body determining healthy aging trajectories. Our findings support the notion that Dlx5/6-dependent regulations in GABAergic interneurons play an important role in the central regulation of behavior and metabolism and ultimately of healthspan and lifespan.

The intimal link between cognitive, rewarding, hypothalamic and peripheral metabolic control systems is most probably at the origin of the comorbidity observed between metabolic syndrome and mental health disorders (64). Individuals with schizophrenia, autism spectrum disorder and other psychiatric conditions have a higher prevalence of metabolic syndrome compared to the general population (65). Reciprocally, obesity impairs cognition and produces atrophy of brain regions associated with learning and memory deficits. Metabolic and psychiatric disorders are both associated with an increased risk of all***-***cause mortality (66). The pathophysiological, molecular and cellular mechanisms linking metabolic and psychiatric disorders to shorter healthspan are still poorly understood. A genetic component is suggested by the fact that several genetic conditions, such as Prader-Willi syndrome (PWS), present at the same time metabolic and cognitive impairment (67) and reduced life expectancy. Interestingly, it has been shown that Dlx5 promotes GABAergic differentiation participating to a protein complex which includes also MAGE-D1 and Necdin (NDN)(36, 37) one of the genes affected in PWS. Moreover, the *DLX5/6* bigenic cluster is close to loci associated to autism (68-71).

A remarkable association between *DLX5/6* and lifespan comes from the observation that these genes are progressively and linearly hypermethylated during human aging (43, 44, 72) and during cellular senescence (45). *DLX5/6* hypermethylation has also been observed in Down syndrome patients with cognitive and metabolic impairment (73). Furthermore the methylation state of individual CpG sites of both *DLX5* and *DLX6* has been shown to increase in adipose tissue and correlate with age and BMI of both female and male cohorts (74).

In the adult, *Dlx5* and *Dlx6* are not only important in the brain, but play also a central role in determining male (75) and female reproduction (50). Our finding that *Dlx5/6* are also important in determining healthspan and lifespan lead us to suggest that they might play an important role in establishing a fecundity/lifespan tradeoff during evolution. In this respect it is important to note that the methylation level of *DLX5/6* increases in response to aging (43), metabolism (74) and life exposures (76) suggesting that these genes might be integrators of lifespan determinants.

## METHODS

Detailed description of methods including: animal procedures, genotypig, histological analysis, single cell RNA sequencing clustering, behavioral tests, coat condition scoring, and statistical tests is provided as Supplementary Information

## ACKNOWLEDGMENTS

This research was partially supported by the EU Consortium HUMAN (EU-FP7-HEALTH-602757) to G.L. and by the ANR grants TARGETBONE (ANR-17-CE14-0024) to GL and METACOGNITION (ANR-17-CE37-0007) to GL, S.L., F.T and an ATM grant to NN-N, CdL is supported by a grant of the French Ministry of Research.

A particular thank goes to the team in charge of mouse animal care and in particular M Stéphane Sosinsky and M Fabien Uridat and to Pr. Amaury de Luze in charge of the “Cuvier” ethical committee. We thank Mlle Rym Aouci, Mlle Mey El Soudany, M Zakaria Maakoul, Mlle Marianne Pungartnik, M Guillaume Robert, Mlle Marjorie Sabourin, and M Benjamin Vanhoutte for experimental support. We thank Mss. Aicha Bennana and Lanto Courcelaud for administrative assistance.

## Author contributions statement

**C.d.L.** Performed experiments, analyzed data, prepared figures, contributed to the text

**E.H.** Performed single cell data analysis

**G.A.** Performed experiments

**A.F.** Performed experiments

**R.H.** Performed body composition analysis

**C.V.** Performed social interaction tests

**F.d.C.** Performed social interaction tests

**C.O-M.** Designed social interaction tests

**S.P.** Performed social interaction test

**F.T.** Provided experimental support, results evaluation and data analysis

**E.E.** Performed social behavior tests

**T.B.** Provided experimental support, results evaluation and data analysis

**S.L.** Provided experimental support, results evaluation and data analysis

**G.L.** Coordinated the work, analyzed data, wrote the text, prepared figures, and took histological images.

**N.N.-N.** Performed and coordinated the experimental work, wrote the text.

### Animal experimentation approvals

Procedures involving animals were conducted in accordance with the directives of the European Community (council directive 86/609), the French Agriculture Ministry (council directive 87–848, 19 October 1987, permissions 00782 to GL) and approved by the “Cuvier” ethical committee.

