## Supplementary Figures and Methods for "*Dlx5* and *Dlx6* expression in GABAergic neurons controls behavior, metabolism, healthy aging and lifespan"

#### Short title: *Dlx5/6* control behavior, metabolism and lifespan.

Camille de Lombares[1], Eglantine Heude[1], Gladys Alfama[1], Anastasia Fontaine[1], Rim Hassouna[2], Cécile Vernochet[3], Fabrice de Chaumont[4], Christophe Olivo-Marin[4], Elodie Ey[5], Sébastien Parnaudeau[3], François Tronche[3], Thomas Bourgeron[5], Serge Luquet[2], Giovanni Levi[1,*,+] and Nicolas Narboux-Nême[1,*]

* Co-senior authors

1. Évolution des Régulations Endocriniennes, CNRS, UMR-7221, Muséum National d'Histoire Naturelle, Département AVIV – 7 rue Cuvier, 75005 Paris, France
2. Unité de Biologie Fonctionnelle et Adaptative (BFA), Université Paris Diderot, Sorbonne Paris Cité, CNRS UMR 8251, 75205, Paris, France
3. Team « Gene Regulation and Adaptive Behaviors », Neurosciences Paris Seine, INSERM U 1130, CNRS UMR 8246, Paris, France.
4. CNRS UMR 3691, BioImage Analysis Unit., Institut Pasteur, Paris, France
5. CNRS UMR 3571, Human Genetics and Cognitive Functions, Institut Pasteur, Paris, France.

\+ Correspondence should be addressed to:

Giovanni Levi
UMR7221 CNRS/MNHN
7, rue Cuvier
75231 Paris, Cedex 05, FRANCE


**Keywords:** Aging, GABAergic neurons, Dlx5, Dlx6, metabolism, behavior, brain

### Detailed Materials and Methods

#### Animals

Procedures involving animals were conducted in accordance with European Community (Council Directive 86/609) and French Agriculture Ministry directives (Council Directive 87–848, permissions 00782 to GL). The project was reviewed and approved by the "Cuvier" ethical committee of the Muséum National d'Histoire Naturelle (approval n° 68-028r1) validated by the French Ministry of Agriculture.

Mice were housed in light, temperature (21°C) and humidity (50–60% relative humidity) controlled conditions. Food and water were available ad libitum. Mice were individually identified by microchip implanted 3 weeks postnatal and litter size and a genotypes were kept in record. WT animals were from Charles River, France. $Dlx5/6^{flox/flox}$ mice (1) were backcrossed and bred on a mixed C57BL6/N X DBA/2N genetic background.

$Slc32a1^{tm2(cre)Lowl}$ knock-in mice (here referred as $Vgat^{cre/+}$ mice) were purchased from Jackson Laboratories through Charles River, France. To obtain the double conditional mutant $Vgat^{\Delta Dlx5\text{-}6/+}$ and $Vgat^{\Delta Dlx5\text{-}6}$ in which the DNA-binding regions of both *Dlx5* and *Dlx6* are deleted by GABAergic-specific cre-recombinase, we crossed $Vgat^{cre/+};Dlx5/6^{flox/+}$ males with $Dlx5/6^{flox/flox}$ females (Fig.S3A). $Dlx5^{lacZ}$ mice (2) were also backcrossed and bred on a mixed C57BL6/N X DBA/2N genetic background.

#### Genotyping

For genotyping, DNA was extracted from mice tails using a KAPA express extraction kit (Kapa Biosystems, Sigma, France). Control, $Vgat^{\Delta Dlx5\text{-}6/+}$ and $Vgat^{\Delta Dlx5\text{-}6}$ mice were genotyped by PCR using allele-specific primers using TAKARA Ex Taq (Takara).

To identify the $Dlx5^{flox}$ and $Dlx5^{\Delta}$ alleles the following three primers were used:

**d** *5′*-TTCCATCCCTAAAGCGAAGAACTTG-*3′*

**e** *5′*-CCTCCCAGAAATACCCCTTCTCTTG-*3′*

**f** *5′* -GTCCCATCCTCAGATCAC -*3′*

Wild-type, $Dlx5^{flox}$ and $Dlx5^{\Delta}$ alleles give rise respectively to 106 bp, 216 bp and 244 bp PCR products.

To identify the $Dlx6^{flox}$ and $Dlx6^{\Delta}$ alleles the following three primers were used:

**a** *5′*-CTTTAGGCGTTGGGAAAGCCAGG-*3′*

**b** *5′*-GCATTATGATAGTGGATCGAATCTAG -*3′*

**c** *5′*-CTGGTCTCAGCTCATAAGTTTCCTTC-*3′*

Wild-type, $Dlx6^{flox}$ and $Dlx6^{\Delta}$ alleles give rise respectively to 165 bp, 222 bp and 345 bp PCR products.

**RT-PCR analysis**

Total RNA was isolated from cortical fragments of control, $Vgat^{\Delta Dlx5\text{-}6/+}$ and $Vgat^{\Delta Dlx5\text{-}6}$ mice using an RNeasy minikit (Qiagen) according the manufacturer instructions. On-column deoxyribonuclease (Qiagen) digestion was incorporated into an RNA isolation procedure to remove potential genomic DNA contamination. RNA concentration and the ratio of the absorbance at 260 and 280 nm were measured using a NanoDrop 2000 spectrophotometer (Thermo Scientific). Reverse transcription was carried out using 600 or 200 ng total RNA and Superscript III (Invitrogen) or Primscript (Ozyme) reverse transcriptase to obtain cDNA.

*Dlx5* and *Dlx6* transcripts were analysed using the following primers (see Fig. S4):

**p4** GTCCCAAGCATCCGATCCG

**p3** CAGGTGGGAATTGATTGAGCTG

**p1** ACATTACCCTCTGCACTGCTTG

**p2** ATGTAGCTGTTGGGAGGCATAC

##### Histological analysis

Mice were fixed by transcardiac perfusion with 4% paraformaldehyde and brains were post-fixed by overnight immersion in 4% paraformaldehyde at 4°C. Samples were cryoprotected in 30% sucrose and frozen. Cryoprotected brains were embedded in OCT (Leica, France) and 60-µm-thick cryostat sections were prepared.

For *lacZ* expression analysis, adult brains were fixed by perfusion with 4% PFA with no postfixation. X-gal staining was performed as described (3). Immunohistochemistry on tissue sections was performed on cryostat sections (60 µm) of adult *Dlx5*$^{lacZ/+}$ brains, incubated overnight at 4°C with a chicken anti β-D-galactosidase antibody (1:2000; Aves labs BGL-1040) combined with either mouse anti PV (1:2000, Sigma P3088), rabbit anti Calretinin (1:1000, Millipore AB5054) or rat anti Somatostatin (1:1000, Millipore MAB354). Sections were then incubated for 2 hours at room temperature in the corresponding secondary fluorescent antibodies (1:300; Jackson Immunoresearch). Pictures were acquired using a Leica SP5 confocal microscope.

##### Single-cell RNA sequencing clustering

Single-cell RNA sequencing (scRNA-seq) analysis was performed on publicly available datasets where individual cells from adult frontal cortex were profiled using a unique molecular identifier protocol (GEO accession number GSE116470) (4). Digital gene expression (DGE) matrices from 21 sequencing pools including a total of 190,972 cells were compiled before principal component-based analysis (PCA).

The R package Seurat was used for cell clustering analysis (5). To ensure quality of data, we excluded cells with fewer than 500 sequenced transcripts, and outliers cells with more than 6000 sequenced transcripts and high mitochondrial percentage (>15%). Thus, 130,845 cells from adult frontal cortex were finally used for clustering analysis. To identify a set of highly variable genes, we first calculated the variance and mean for each gene in the dataset and sorted genes by their variance/mean ratio. We then selected the top 1,000 highly variable genes for downstream PCA. These selected genes were then centered and scaled across all cells and PCA was performed with 60 components. The analysis resulted in 42 clusters across the global frontal cortex.

To identify finer substructures among *Dlx5*-positive single cells, the subset was selected and subjected to another round of variable gene selection, scaling and PCA with 20 components. A total of 967 *Dlx5*-positive single cells distributed in 7 clusters were identified.

**Behavioral tests**

Mice were taken to the test room 30 min before the test. Behavioral procedures were conducted between 10 a.m. and 4 p.m. in a dim and quiet room. Observers were blind to the experimental design.

***Open field test (OFT)***

The OFT is widely applied to test motor and anxiety-like behaviors of rodents in a novel environment (6 , 7). The equipment consisted of a close square arena (72 × 72 cm). The computer defined the grid lines dividing the box floor into 16 equal-sized squares, with the central four squares regarded as the center. Each mouse aged of less than one year, was gently placed at one corner of the arena facing the wall and video taped for 10 min. All animals were tracked and recorded with a digital camera and analysed by Ethovision system (Noldus). Delay to enter in the center, time in the center, number of entries into the center, total distance covered, average and peak velocity and acceleration were analysed. After each test, the equipment was cleaned and disinfected.

***Marble burying test (MBT)***

The marble burying test (MBT) (8) was employed to measure anxiety and compulsive-like behaviors. A clear Plexiglas box (36,5 cm long × 20,7 cm wide × 14 cm high) was filled to a depth of 3 cm with standard wood shavings. Twenty glass marbles were placed on the surface of the shavings. Mice aged of less than 6 months were individually placed in the center of the box; the test session was 10 min. At the end of the session, a picture of the marbles was taken, and the marbles buried index was counted with the Fiji (ImageJ) image-processing program.

***Nest building test***

The nest-building test was performed as described (9). Each mouse aged of less than one year, was housed in a single cage before testing. During the test, a paper towel (30 cm × 21,5 cm) was placed in the cage and left for one week. The quality of the nest was scored each following day into four categories: perfect nest built, paper completely torn, paper partially torn and no interaction with paper. Nests were scored at 10h a.m.

***Sociability tests***

*1) Social interaction test*

Social interaction test was performed as described in Barik et al. (10). Briefly, mice (aged 5-6 months) were introduced for 150 s in an open-field (40cm x 40cm x 25) containing an empty perforated polycarbonate box (« no target » condition). Immediately after this, mice were rapidly removed and an unfamiliar male mouse was placed in the box (« target » condition) and mice were re-exposed to the open-field for another 150 s. The time spent in the interaction zone surrounding the polycarbonate box while empty or with an unfamiliar mouse was recorded and used as an index of social interaction.

*2) Open field social behavior*

After one hour of habituation a female control mouse was placed in a test cage (50x50x30 cm) together with either a second control, a *Vgat*$^{\Delta Dlx5\text{-}/+}$ or a *Vgat*$^{\Delta Dlx5\text{-}6}$ female mouse. Mice were then filmed for 10 min. Social behavior was measured using real-time approach that couples computer vision, machine learning and Triggered-RFID identification to track and monitor animals (11).

***Locomotor activity***

Locomotor activity of the mice (aged 2-5 months) was measured for 90 min for 3 consecutive days as described (12). Mice were introduced in circular chambers (4.5-cm width, 17-cm external diameter) crossed by four infrared captors (1.5 cm above the base) placed at every 90° (Imetronic, Bordeaux, France). The locomotor activity was counted when animals interrupted two successive beams and thus had travelled a quarter of the circular corridor and was expressed as ¼ turns per 90 min.

**Scoring of coat conditions**

Groups of mice (n=12 each) were individually photographed and observed at different ages. The parameters measured were: A) alopecia level scored into three grades: score 1: alopecia on all body or at least two zones; score 0.5: alopecia on one body zone; and 0: no alopecia observed; B) loss of fur color also scored into three grades: score 1: loss of fur color on all body or at least two zones; score 0.5: loss of fur color on one body zone; and 0: no loss of fur color observed; C) loss of whiskers scored as follows: score 1: complete loss of whiskers; score 0.5: partial loss of whiskers; and 0: no loss of whiskers observed, and finally; D) coat conditions scored as follows: 1: ungroomed, ruffled, non-shiny appearance; 0,5: average appearance; 0: smooth, shiny coat. The global scoring of coat condition was performed as described (13).

##### Statistical analyses

The Pearson's chi-squared, ANOVA and Kruskal-Wallis test were conducted using Prism (Graphpad Software, La Jolla, CA, USA) to calculate the differences between groups.

All values are expressed as means ± SEM of combined data from replicate experiments. Values of $P < 0.05$ were considered statistically significant.

### SUPPLEMENTARY FIGURES

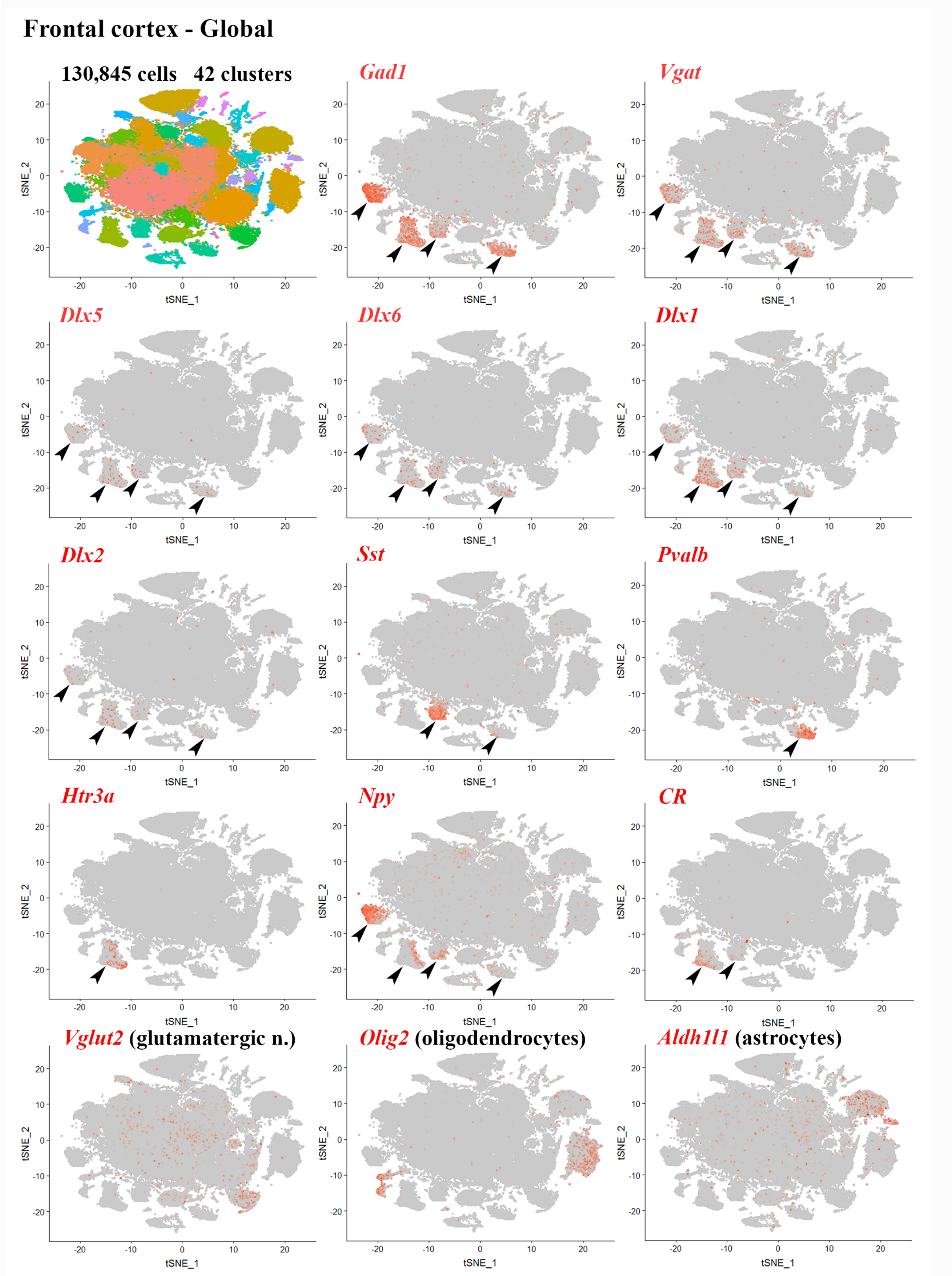


**Supplementary Figure 1) Frontal cortex single-cell clustering and expression of markers in distinct subclusters.**

(Upper-left panel) t-distributed stochastic neighbor embedding (t-SNE) plot showing the overall gene expression relationship among 130,845 single cells isolated from the frontal cortex (4) using principal component analysis (PCA). The 42 different cell clusters are color-coded.

(Other panels) t-SNE plots showing expression of markers for cortical GABAergic neurons (*Gad1, Vgat, Dlx5, Dlx6, Dlx1, Dlx2, Sst, Pvalb, Htr3a, Npy, CR*), Glutamatergic neurons (*Vglut2*), Astrocytes (*Aldh1l1*) and Oligodendrocytes (*Olig2*). *Dlx5/Dlx6*-positive cells are present in all GABAergic subclusters (arrowheads), but not in glutamatergic neurons, oligodendrocytes and astrocytes.

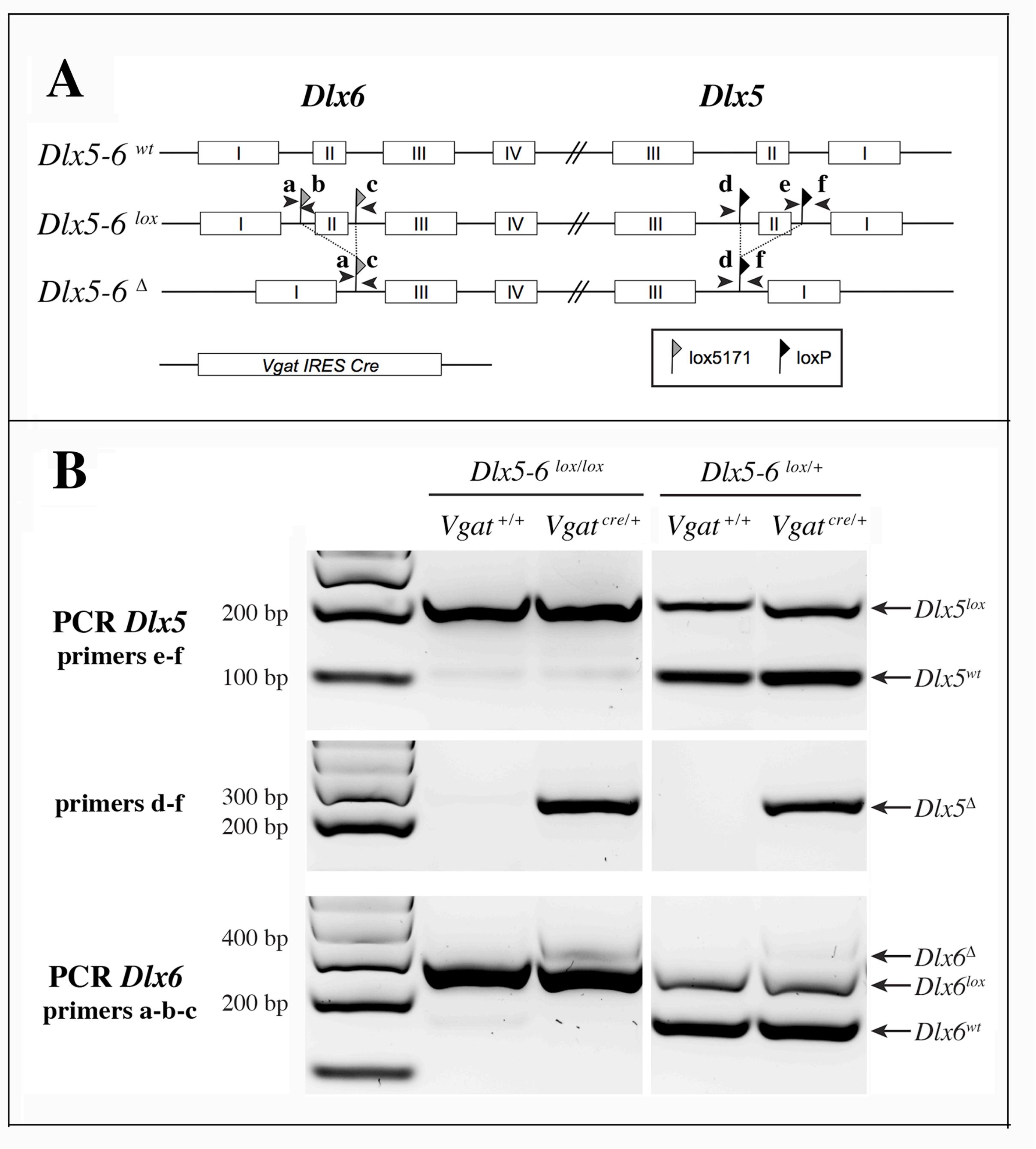


**Supplementary Figure 2) Strategy of *Dlx5* and *Dlx6* simultaneous invalidation and mouse genotyping.**

A) Exons 2 of *Dlx5* and *Dlx6* were respectively framed with *loxP* and *lox5171* sequences as described in (1). In the presence of a *Slc32a1-IRES-Cre (Vgat-Cre)*, exons II of

*Dlx5* and *Dlx6* are deleted in GABAergic interneurons generating a *Dlx5-6*Δ allele. Arrowheads indicate the position and name (a to f) of the primers used for genotyping.

B) PCRs on cortical DNA extracts. Primers a, b, c and d, e, f were respectively utilized to reveal *Dlx6* and *Dlx5* recombination. The floxed and wild type *Dlx5* alleles (primers e-f) were revealed in a separate PCR than that used to reveal the recombinant (*Dlx5Δ*) allele (primers d-f). Wild type, floxed and recombinant *Dlx6* alleles were identified with a single PCR with primers a, b, c. In the presence of Cre recombinase, a band corresponding to the recombinant allele (Δ) can be detected for both *Dlx5* (primers d-f) and *Dlx6* (primers a, b, c) in $Vgat^{\Delta Dlx5\text{-}6}$. In $Vgat^{\Delta Dlx5\text{-}6/+}$ mice wild type and recombinant alleles are detected.

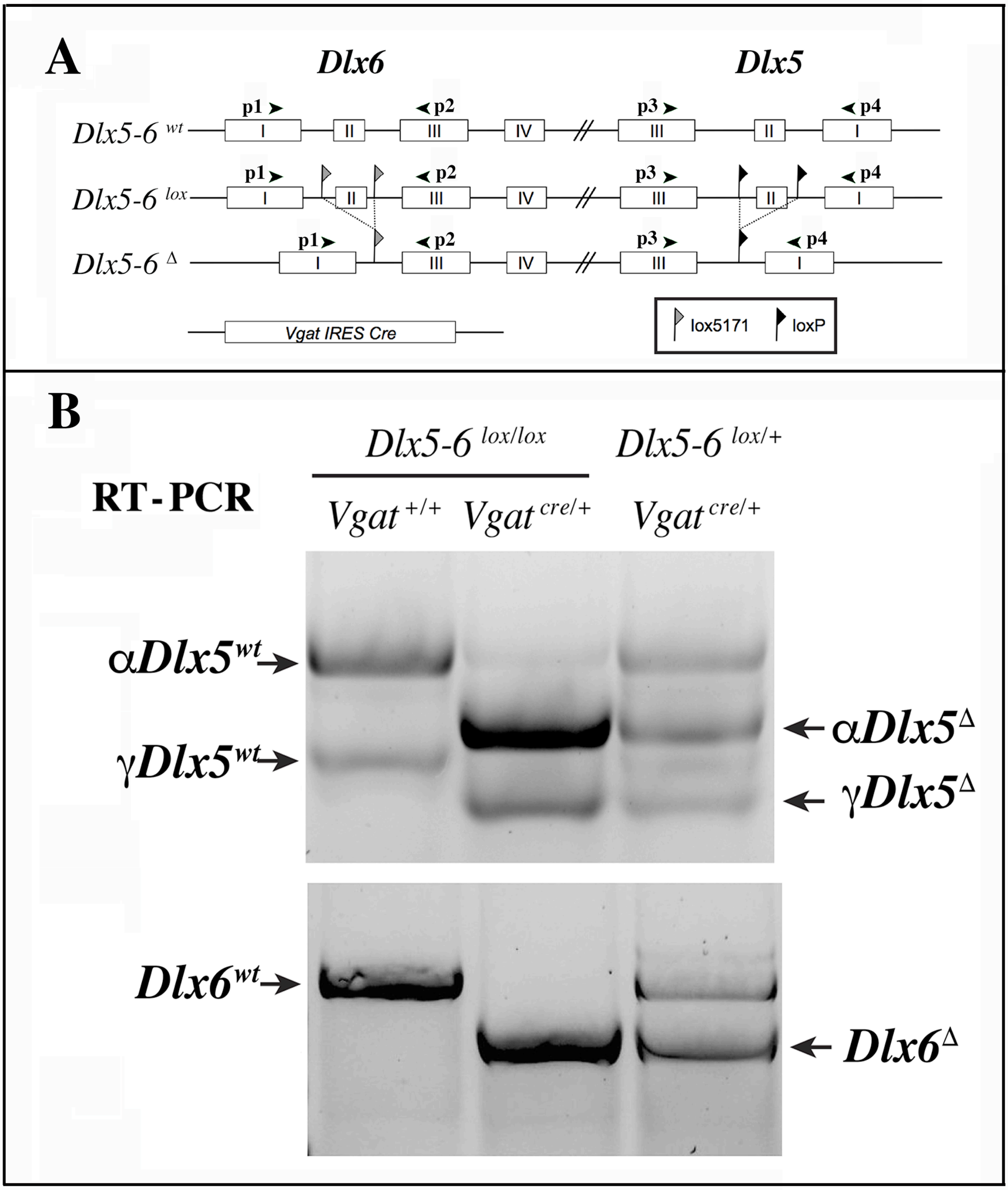


**Supplementary Figure 3) RT-PCR analysis of *Dlx5* and *Dlx6* expression in the cerebral cortex**

A) Primers p1 to p4 were used to analyze the presence of *Dlx5* and *Dlx6* transcripts in reverse-transcribed RNA extracts from adult cerebral cortex fragments.

B) Two known splice variants of *Dlx5 (αDlx5* and *γDlx5*, (14)) were amplified with primers p3 and p4. Deletion of exon II shifted both bands giving rise to *αDlx5*$^{\Delta}$ and *γDlx5*$^{\Delta}$. A small fraction of *Dlx5* transcripts, possibly corresponding to expression of this gene in non-GABAergic cells, was not recombined. *Dlx6* transcripts were amplified with primers p1 and p2 and did not show any splice variants.

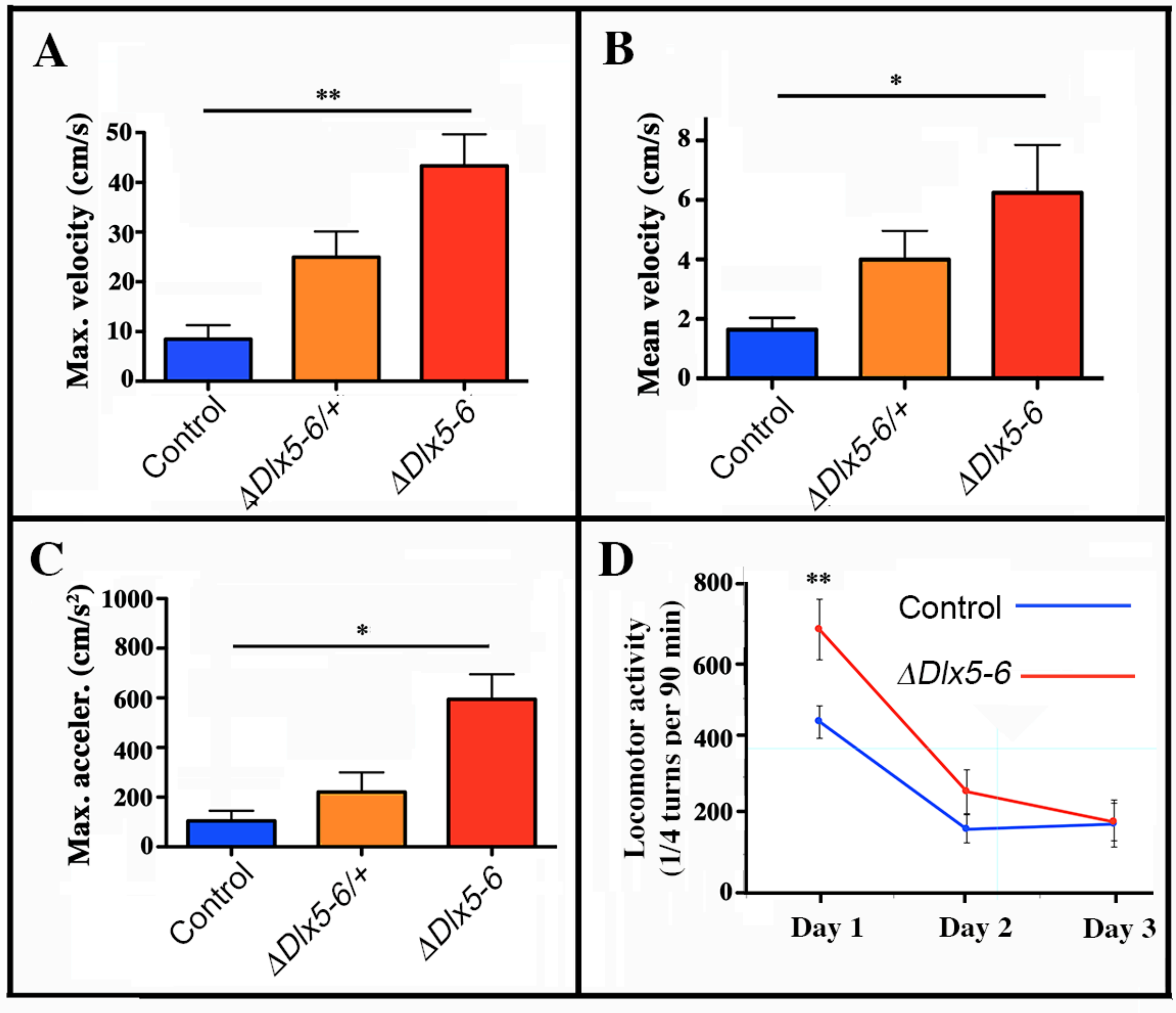


**Supplementary Figure 4) Mouse velocity and acceleration in an open field test**

Maximal (A) and average (B) velocity and maximal acceleration (C) of mice in the open field test (see Figure 2). During the 10 minutes test the velocity and acceleration of mutant mice was significantly higher than that of controls. D) Locomotor activity was measured on a 90 min period in a circular corridor for three consecutive days. Only the first day a significant difference was observed suggesting an increased response to novelty of mutant mice. One way (A, B, C) and repeated two way (D) ANOVA tests, Bonferroni post-hoc test $p<0.01$ were performed. Histograms bars indicate the mean±SEM. **: $p<0.01$; *: $p<0.05$.

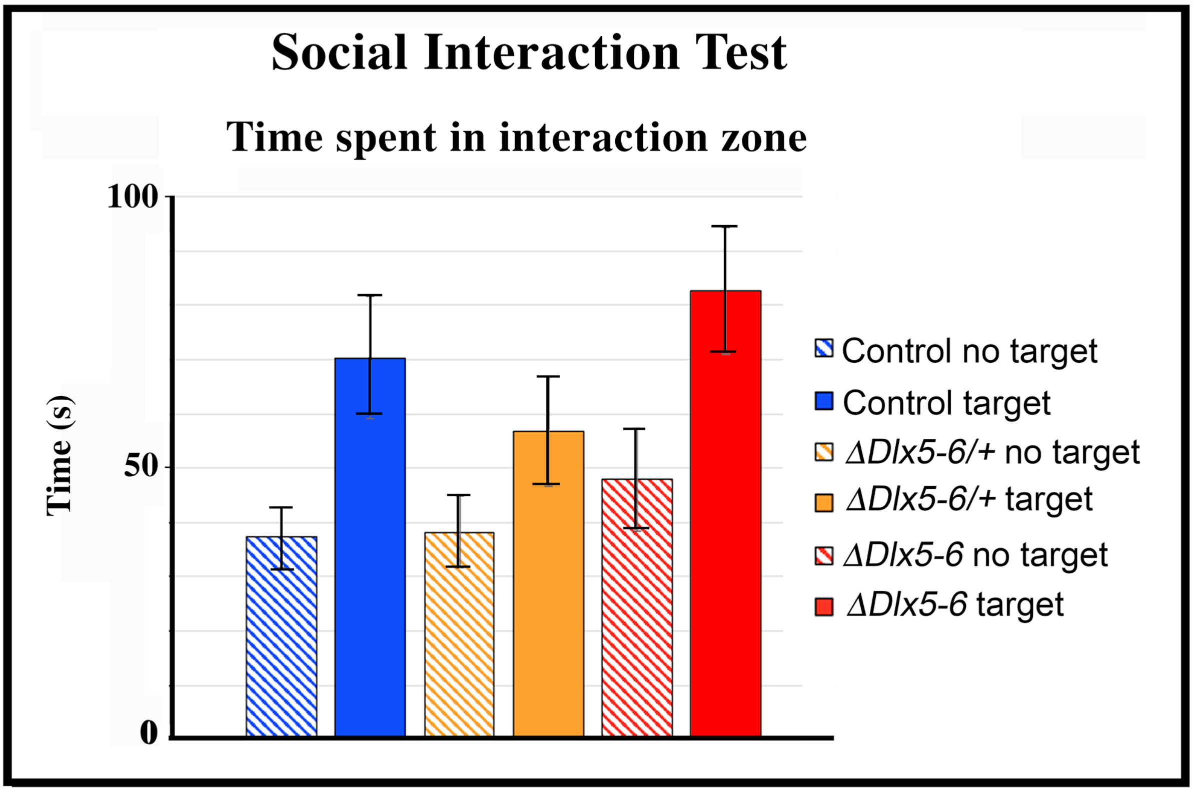


**Supplementary Figure 5) Social interaction sociability tests**

The time spent establishing social contacts with an unfamiliar mouse confined (target) in a transparent and perforated box placed in the center of an open field was compared to the time spent in proximity of the same empty box (no target). Independently from their genotype, mice spent more time in the interaction zone when a target mouse of the same sex was present in the box (controls: 37±5 s "no target" vs 70±11 s "target"; $Vgat^{\Delta Dlx5\text{-}6/+}$: 38±5 s "no target" vs 57±10 s "target"; $Vgat^{\Delta Dlx5\text{-}6}$: 48±9 s "no target" vs 83±12 s "target"). Control, $Vgat^{\Delta Dlx5\text{-}/+}$ and $Vgat^{\Delta Dlx5\text{-}6}$ mice did not show any significant difference in the time spent interacting with the unfamiliar mouse (nor in the time spent interacting with the empty box), suggesting that the genotype does not impact on the social behavior (two-way repeated ANOVA "no target" vs "target").

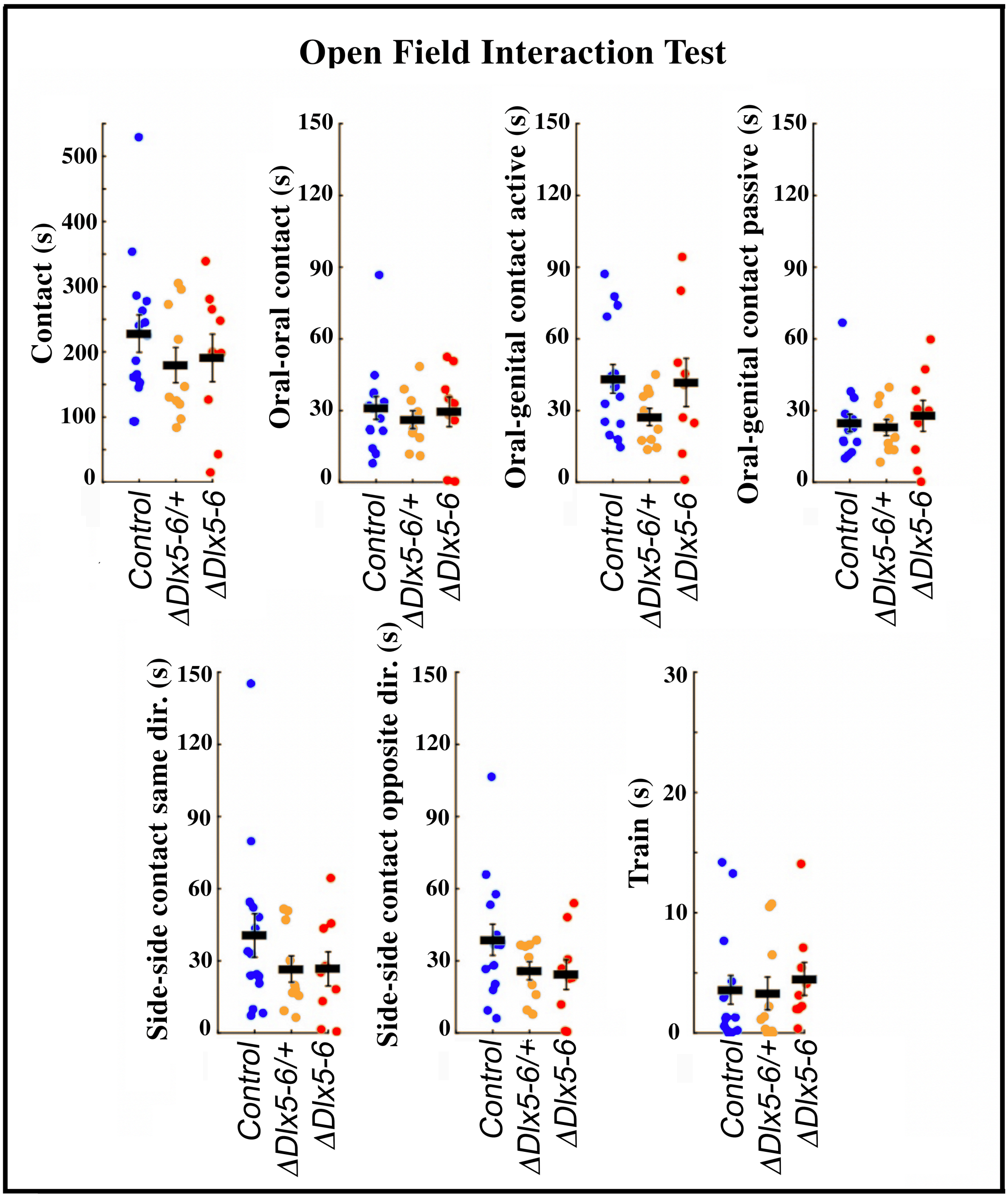


**Supplementary Figure 6) Open field social behavior**

The social behavior of couples of mice placed simultaneously in an open field was measured using a real-time procedure that couples computer vision, machine learning and Triggered-RFID identification to track and monitor animals (11). The system extracts a thorough list of individual and collective behavioral traits and provides a unique phenotypic profile for each animal. None of the analyzed socialization parameters showed any significant difference by Wilcoxon test.

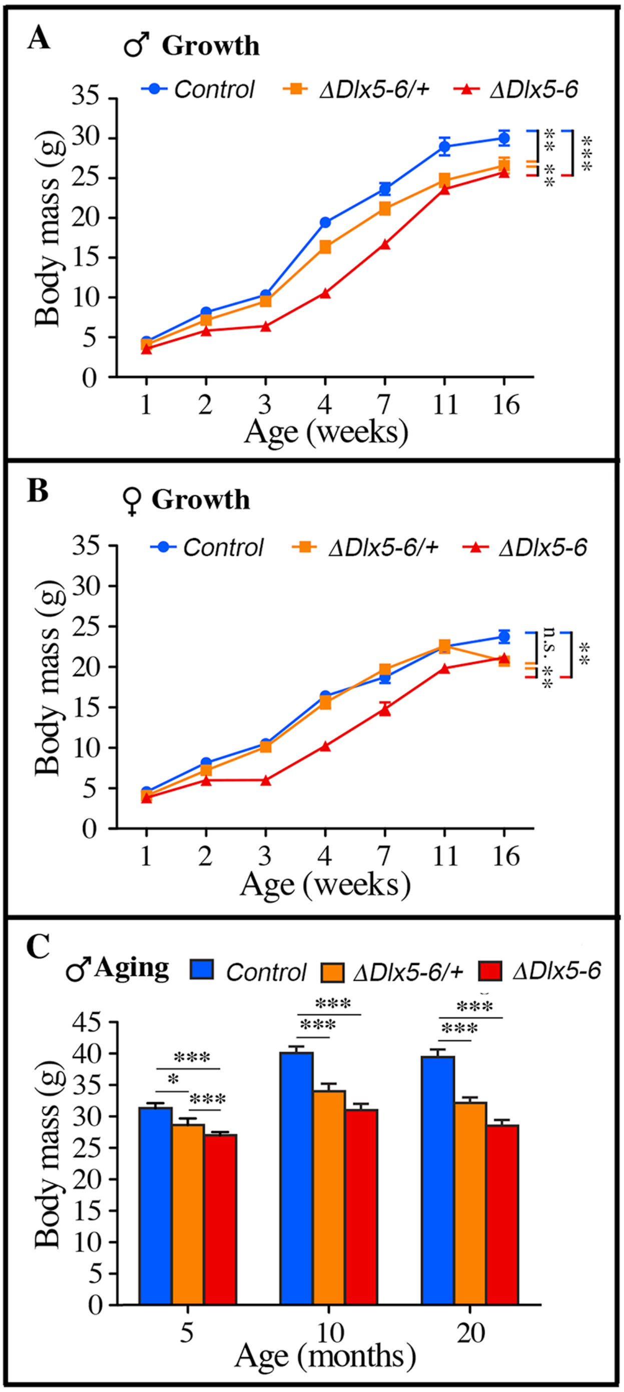


**Supplementary Figure 7) Weight measures during growth and aging.**

A-B) The body weight of cohorts of male (A) and female (B) control, *Vgat*$^{\Delta Dlx5\text{-}6/+}$ and *Vgat*$^{\Delta Dlx5\text{-}6}$ mice was measured during the first 16 weeks of growth. At all time points, *Vgat*$^{\Delta Dlx5\text{-}6}$ mice (male and female) displayed a significant weight reduction; *Vgat*$^{\Delta Dlx5\text{-}6/+}$

males had also a significantly lower weight, while, during growth, until 11 weeks of age, the weight of female *Vgat*$^{\Delta Dlx5\text{-}6/+}$ was not significantly different than controls (B).

(C) The body weight of a cohorts of male control, *Vgat*$^{\Delta Dlx5\text{-}6/+}$ and *Vgat*$^{\Delta Dlx5\text{-}6}$ mice (n ≥ 8 per group) was measured during the first 20 months of aging. At all time points analyzed *Vgat*$^{\Delta Dlx5\text{-}6/+}$ and *Vgat*$^{\Delta Dlx5\text{-}6}$ male mice presented a highly significant weight reduction.
